# Discovery and characterisation of novel circular bacteriocin plantacyclin B21AG from *Lactobacillus plantarum* B21

**DOI:** 10.1101/2020.04.10.035774

**Authors:** Aida Golneshin, Mian Chee Gor, Ben Vezina, Nicholas Williamson, Thi Thu Hao Van, Bee K. May, Andrew T Smith

## Abstract

*Lactobacillus plantarum* B21 isolated from Vietnamese sausage (*nem chua*) has previously displayed broad antimicrobial activity against gram positive bacteria including foodborne pathogens *Listeria monocytogenes* and *Clostridium perfringens*. This study successfully identified the antimicrobial agent as plantacyclin B21AG, a 5668 Da circular bacteriocin demonstrating high thermostability, resistance to a wide range of pH, proteolytic resistance and temporal stability. We report a reverse genetics approach used to identify and characterise plantacyclin B21AG. The bacteriocin was purified from culture supernatant by a short process consisting of concentration, n-butanol extraction and cation exchange chromatography. A *de novo* peptide sequencing using LC-MS/MS techniques identified two putative peptide fragments which were mapped to the genome of *Lactobacillus plantarum* B21. This revealed an ORF corresponding to a putative circular bacteriocin with a 33-amino acid leader peptide and 58-amino acid mature peptide found on native plasmid pB21AG01. The corresponding gene cluster, consisted of seven genes associated with post-translational circularisation, immunity and secretion. The robust nature of plantacyclin B21AG, its antimicrobial activity and associated machinery for cyclisation make it an interesting biotechnological target for further development, and application as a food-safe antimicrobial.

## Introduction

Food borne disease due to pathogen contamination is a major concern in the food industry [1]. Reports estimate that 4.1 million foodborne gastroenteritis cases occurs in Australia annually, costing the country about $1.2 billion per year [2, 3]. Discovery of antimicrobial agents which contribute to food safety will have economic and social significance and are a top priority of the food industry. There are building health concerns regarding current chemical food preservatives and additives used in food [4], which has resulted in interest and development of foods without chemical additives. The increased awareness of food safety amongst consumers has created demand for natural products such as those produced by bacteria, to control the growth of food spoilage bacteria and food-borne pathogens [5].

Lactic acid bacteria (LAB) have been long associated with food and feed fermentation, specifically as a starter culture in the production of fermented meat, vegetables, fruits, alcoholic beverages, dairy products and silage [6, 7]. They have been shown to inhibit the growth of food spoilage bacteria, as well as have a beneficial influence on nutrition, organoleptic and shelf-life characteristics of fermented products [8–10]. The preservative effect of LAB is due to the production of antimicrobial substances, including direct competition, organic acids, hydrogen peroxide, diacetyl, bacteriocins and bacteriocin-like inhibitory substances [10, 11]. Among these antimicrobial substances, bacteriocins have received particular attention in recent years because of their application as natural preservatives in the food industry, and as potential antibiotics targeting multi-drug resistance pathogens [5, 12].

Over the past few decades, many bacteriocins have been identified and studied extensively in LAB [13]. Several approaches have been taken to classify bacteriocins. The original classification of bacteriocins from LAB was suggested by Klaenhammer (14) based on the biochemical and genetic properties. Circular bacteriocins are uniquely distinguished from other classes of bacteriocins as they go through post-translational modification, specifically an N to C termini ligation through an amide bond. Circularisation gives them unique characteristics including high thermostability, resistance to a wide range of pH, proteolytic resistance and temporal stability. To date, There are 18 circular bacteriocins being reported to date, including aureocyclicin 4185 [15], enterocin NKR-5-3B [16], amylocyclicin [17], amylocyclicin CMW1 [18], enterocin AS-48 [19], bacA [20], carnocyclin A [21], circularin A [22], thermocin 458 [23], garvicin ML [24], lactocyclicin Q [25], leucocyclicin Q [26], uberolysin [27], acidocin B [28], butyrivibriocin AR10 [29], paracyclicin [30], gassericin A/reutericin 6 [31] and plantaricyclin A [32]. This group of bacteriocins are unique due to their pH and thermal stability, and the ability to resist many proteolytic enzyme activities, making them an interesting target for industry applications [33].

Several *Lactobacillus* spp. were previously isolated from Vietnamese fermented sausage, *nem chua* [34]. Among the isolates, *L. plantarum* B21 demonstrated the strongest antimicrobial activity against a range of bacterial strains due to acid and bacteriocin production [35]. In this study, we purified and characterised a novel bacteriocin from *L. plantarum* B21. We also extended the findings and used a reverse genetics approach to identify the genetic cluster encoding for proteins involved in the production, immunity and transport of the bacteriocin.

## Results and discussion

### Antimicrobial spectrum of the bacteriocin and sensitivity to proteolytic enzymes

The CFS from *L. plantarum* B21 demonstrated a wide antimicrobial spectrum against the LAB strains tested and foodborne pathogens *Clostridium perfringens* and *Listeria monocytogenes* (Table 1). The strongest antimicrobial activity was against closely-related *Lactobacillus* species. In agreement to most of the LAB bacteriocins reported to date, *L. plantarum* B21 was not shown to be active against any gram-negative bacteria tested in this study. Some studies have shown that treatment with EDTA and lysozyme could render the gram-negative bacteria susceptible to LAB bacteriocin, however this was not looked at [13]. The antimicrobial substance(s) produced by this strain had previously shown interesting characteristics such as wide pH tolerance (3 to 10) and thermo-stability (up to 90°C for 20 min) [35]. The proteinaceous nature of the *L. plantarum* B21 antimicrobial activity found in the CFS was confirmed according to Omar et al. [50]. The killing activity of *L. plantarum* B21 CFS showed sensitivity of different levels to proteinase K, trypsin and pepsin (Fig 1). This indicated that the antimicrobial activity of the CFS from *L. plantarum* B21 was due to a protein and not a secondary metabolite. The bacteriocin activity was completely eliminated with proteinase K but only partly eliminated by trypsin and pepsin, which is an unusual characteristic for bacteriocins [41, 43, 50].

**Fig 1.**
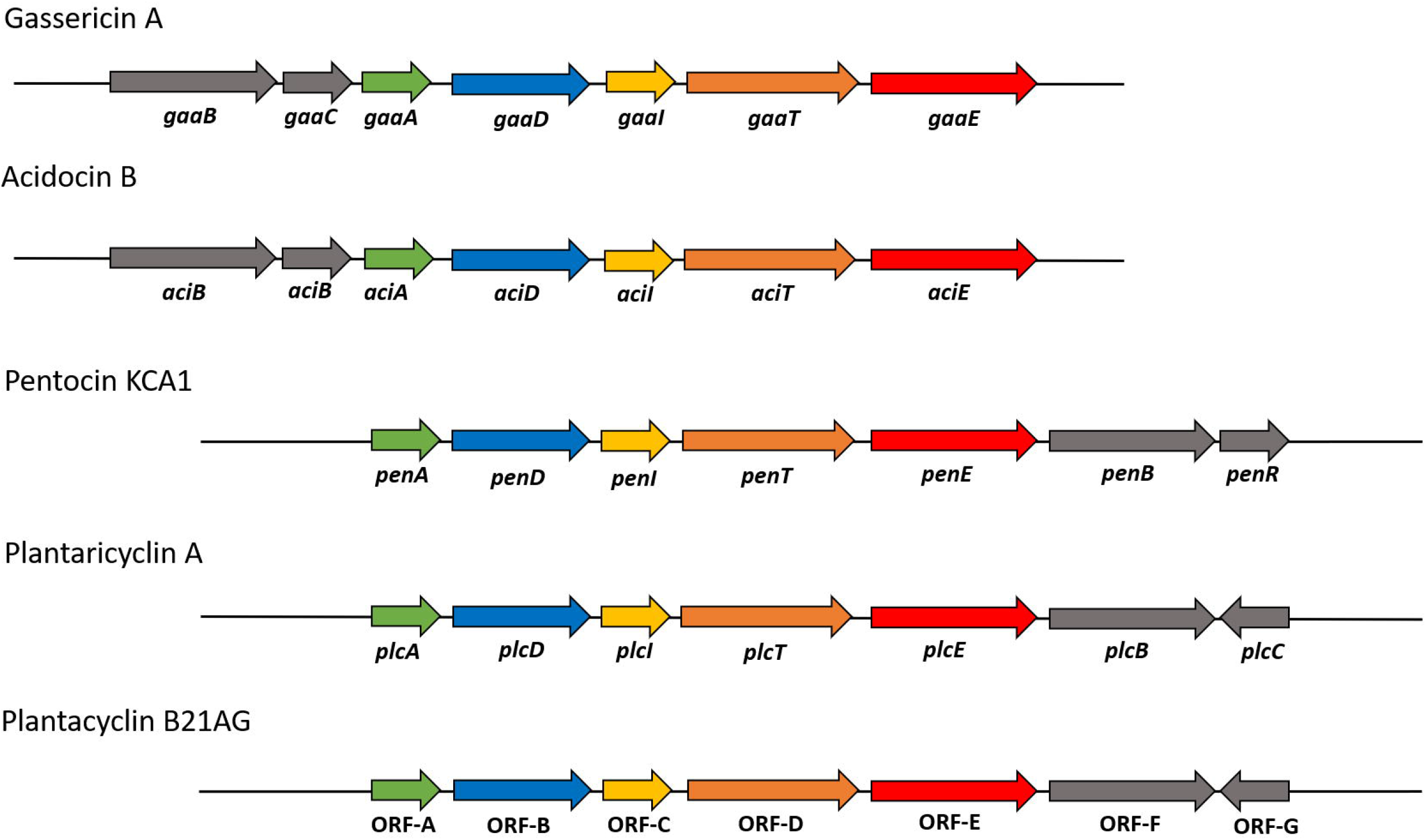
Sensitivity of *Lactobacillus plantarum* B21 bacteriocin to proteolytic enzymes. (1) Proteinase K; (2) Trypsin; (3) Pepsin; (4) Catalase; (5) Untreated cell free supernatant as control.

**Table 1.** Evaluation of *Lactobacillus plantarum* B21 antimicrobial activity by WDA assay

| Indicator strain | Antimicrobial activity* |
| --- | --- |
| *Lactobacillus plantarum* A6 | + |
| *Lactobacillus plantarum* ATCC 8014<sup>a</sup> | + |
| *Lactobacillus arabinosus* 17-5<sup>b</sup> | + |
| *Lactococcus lactis* 345-18<sup>b</sup> | + |
| *Lactobacillus brevis* 19012<sup>b</sup> | + |
| *Lactobacillus casei* ATCC 7469<sup>a</sup> | - |
| *Staphylococcus aureus* ATCC 25923<sup>a</sup> | - |
| *Escherichia coli* ATCC 25922<sup>a</sup> | - |
| *Escherichia coli* ATCC 35218<sup>a</sup> | - |
| *Escherichia coli* NCTC 9001<sup>c</sup> | - |
| *Listeria monocytogenes* 192/1-2 ACM 3173<sup>b</sup> | + |
| *Clostridium perfringens* 52/6-1<sup>b</sup> | + |

\*Positive inhibitory activity [+], No inhibitory activity [-]
*Lactobacillus plantarum* A6 was isolated from nem chua [38]
<sup>a</sup>Strains obtained from American Type Culture Collection (ATCC)
<sup>b</sup>Strains obtained from RMIT University Culture Collection
<sup>c</sup>Strain obtained from National Collection of Type Cultures (NCTC)

### Purification of the bacteriocin

The bacteriocin was purified from the CFS of *L. plantarum* B21 through a series of concentration, solvent extraction, desalting and cation exchange chromatography steps. WDA analysis was performed after every step to monitor the presence of the antimicrobial peptide. The bacteriocin activity was calculated as 800 (AU/mL) against *L. plantarum* A6. The concentration of the *L. plantarum* B21 CFS using a 10 kDa cut off membrane resulted in 16-fold concentration of the bacteriocin, increasing the inhibitory activity from 800 (AU/mL) to 12,800 (AU/mL), while no inhibitory activity was detected in the filtrate (Fig 2a). The concentrated bacteriocin was extracted into n-butanol and the organic phase was freeze dried. The WDA assay of the dried n-butanol phase (resuspended in 20 mM sodium phosphate buffer, pH 6) showed strong bacteriocin activity against *L. plantarum* A6 while no activity was observed in the aqueous phase, confirming n-butanol extraction exhibited complete recovery of the bacteriocin activity (Fig 2b). The extraction and extent of recovery seemed to be even more efficient than other bacteriocins reported where some residual activity is always observed in aqueous phase [41, 51]. The size exclusion and desalting of the bacteriocin using the NAP10 desalting column pre-packed with Sephadex G-25 resin served to help remove excessive lipopolysaccharides and any residual salt in preparation for further chromatography. The FPLC cation exchange chromatography revealed a single peak in absorbance at 214 nm and a small peak at 280 nm corresponding to eluted protein (Fig 2c). The former arising from the peptide bonds and the latter from the aromatic residues of the bacteriocin peptide. The presence of one major peak was due to the selectivity of the n-butanol extraction, combined with the identified bacteriocin was strongly basic, exhibiting an overall positive charge at pH 6.0. Three fractions taken across the single absorbance peak at 214 nm and 280 nm (F14, F15 and E15) showed the strongest bacteriocin activity (Fig 2c). Fractions F14 and F15 were pooled and concentrated, resulting in a yield of approximately 1 mg/L. The yield is considerably lower than those reported in the literature (~2.5 mg/L – 5.5 mg/L) [52, 53]. We speculate about 20-30% of the protein did not bind to the cation exchange column as mass spectrometry indicated the same bacteriocin polypeptide mass in the flow through (data not shown). Slowing the column flow rate and reducing the loading did not improve the yield. One possible explanation could be that a proportion of the bacteriocin was bound to another protein or is highly aggregated, masking its charge properties in some way.

**Fig 2.**
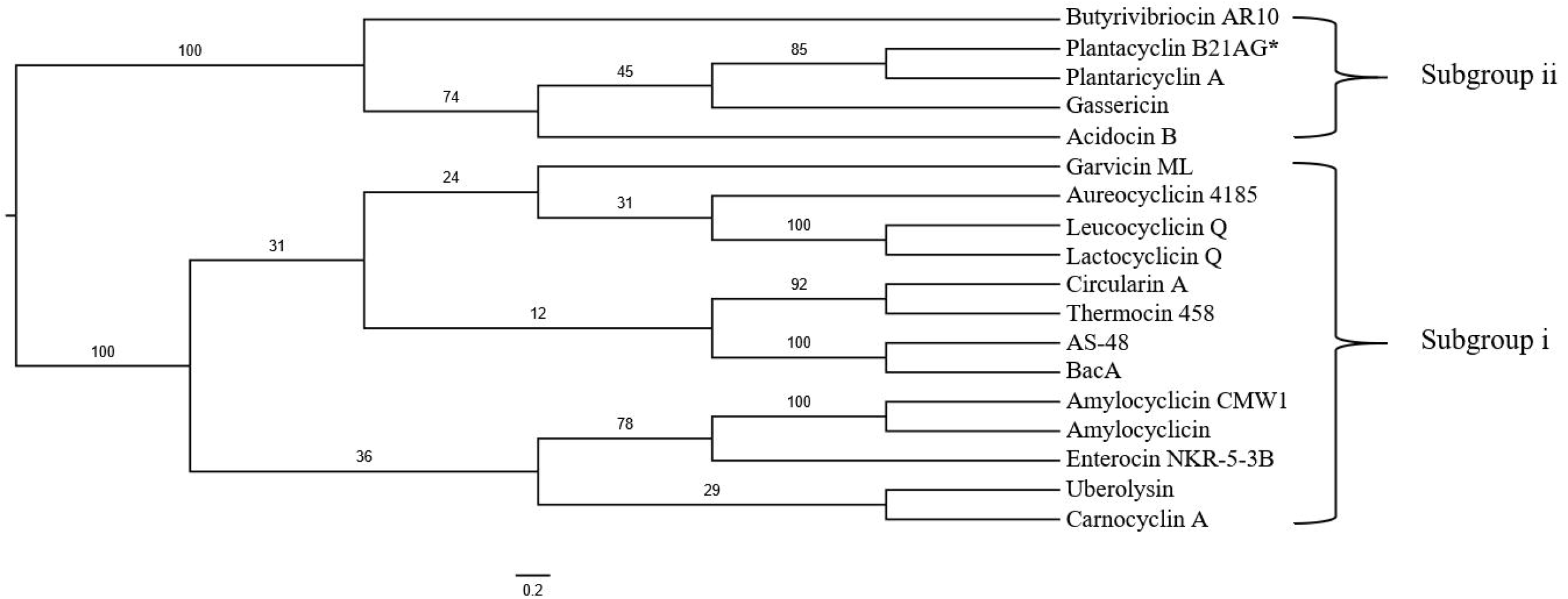
Purification of bacteriocin from the CFS of *Lactobacillus plantarum* B21. *L. plantarum* A6 was used as indicator strain in all WDA assays. (a) *L. plantarum* B21 CFS bacteriocin activity using WDA assay before (left) 800 AU/mL and after (right) at 12,800 AU/mL concentration through an ultrafiltration cell with 10 kDa membrane cut off. (b) The WDA assay examining the antimicrobial activity of organic (left) and aqueous phase (right) from the n-butanol extraction. (c) FPLC of the active fractions from the NAP10 gel filtration column. Eluted protein was detected at 280 nm (blue line) and 214 nm (red line). Fractions F14, F15 and E15 showed the strongest antimicrobial activities based on the WDA assay.

### Molecular weight and purity analysis

The purified bacteriocin showed a single band of approximately 5 kDa on the Tris-Tricine-SDS-PAGE gel (Fig 3a), indicating the purity of the bacteriocin sample. Our result is in accordance to the literature, where a large number of LAB bacteriocin have been reported to be low molecular weight proteins of less than 10 kDa [43, 54–58]. An inhibitory zone was observed around the protein band on the second half of the gel, confirming that the single protein band is responsible for bacteriocin activity (Fig 3b). The mass obtained by MALDI-TOF MS agreed with Tris-Tricine-SDS-PAGE analysis, where one major peak (5668 Da) was observed in the mass spectroscopy analysis (Fig 3c). The second small peak (2830 Da) appeared to be a 2+ species. The MALDI-TOF MS results also showed the presence of the same bacteriocin peptide in the flow through fractions (data not shown). This suggests reduced binding to the cation exchange column either because the protein binding capacity of the column had been exceeded or because the bacteriocin was bound to other proteins. The same phenomenon occurred when the total protein loading on the column was reduced; strongly suggesting either an aggregated state or that it was bound to other proteins.

**Fig 3.**
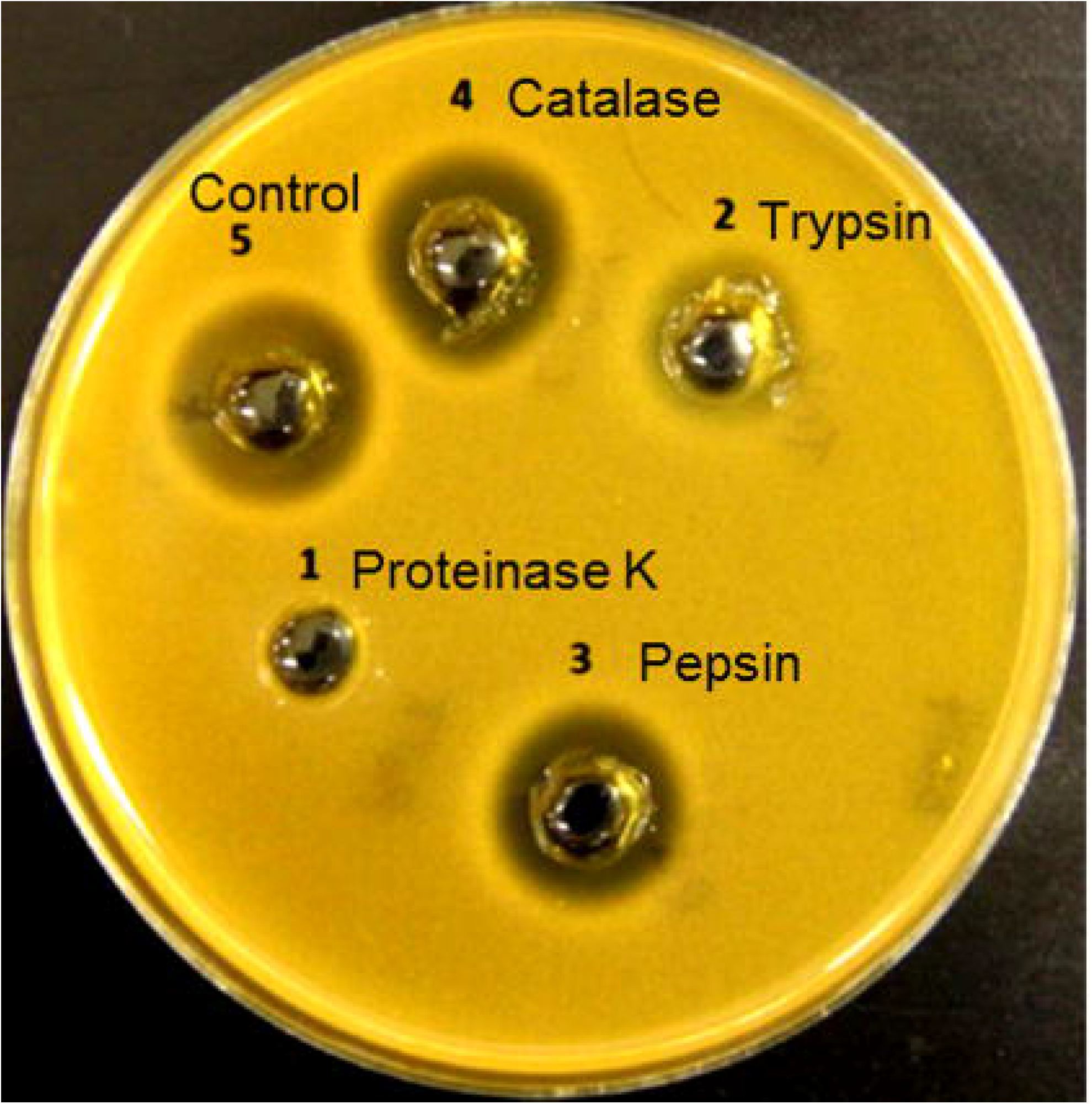
Molecular weight and purity determination by Tris-Tricine-SDS-PAGE gel and MALDI-TOF-MS analysis. (a) SDS-PAGE gel image; lane 1: Low molecular protein marker; lane 2: *L. plantarum* B21 purified bacteriocin. (b) Antimicrobial assay of the SDS-PAGE gel overlayed with the indicator *Lactobacillus plantarum* A6. (c) MALDI-TOF-MS spectrum of the purified *L. plantarum* B21 bacteriocin showing the singly (5668 Da) and doubly (2830 Da) charged species.

When the purified bacteriocin was injected onto a ZORBAX Eclipse analytical C_18_ column (4.6×150 mm) surprisingly, no protein was eluted using an ACN-water gradient up to 60% (ACN) and no peak observed at A_220_ and A_280_. When the ACN concentration gradient was increased to 95%, the bacteriocin protein was successfully eluted from the column using a 40-min linear water-ACN gradient, demonstrated by the presence of a major peak at 37.5 min, corresponding to the *L. plantarum* B21 purified bacteriocin (Fig 4). This experiment was repeated four times and the peaks from all runs overlapped. Failure to elute the bacteriocin protein from a C_18_ column by 60% water-ACN strongly suggested that the *L. plantarum* B21 bacteriocin protein molecule was more hydrophobic than classical bacteriocins. SDS-PAGE electrophoresis, MALDI-TOF-MS and RP-HPLC analyses all confirmed a high level of purity and homogeneity of the final *L. plantarum* B21 bacteriocin sample.

**Fig 4.**
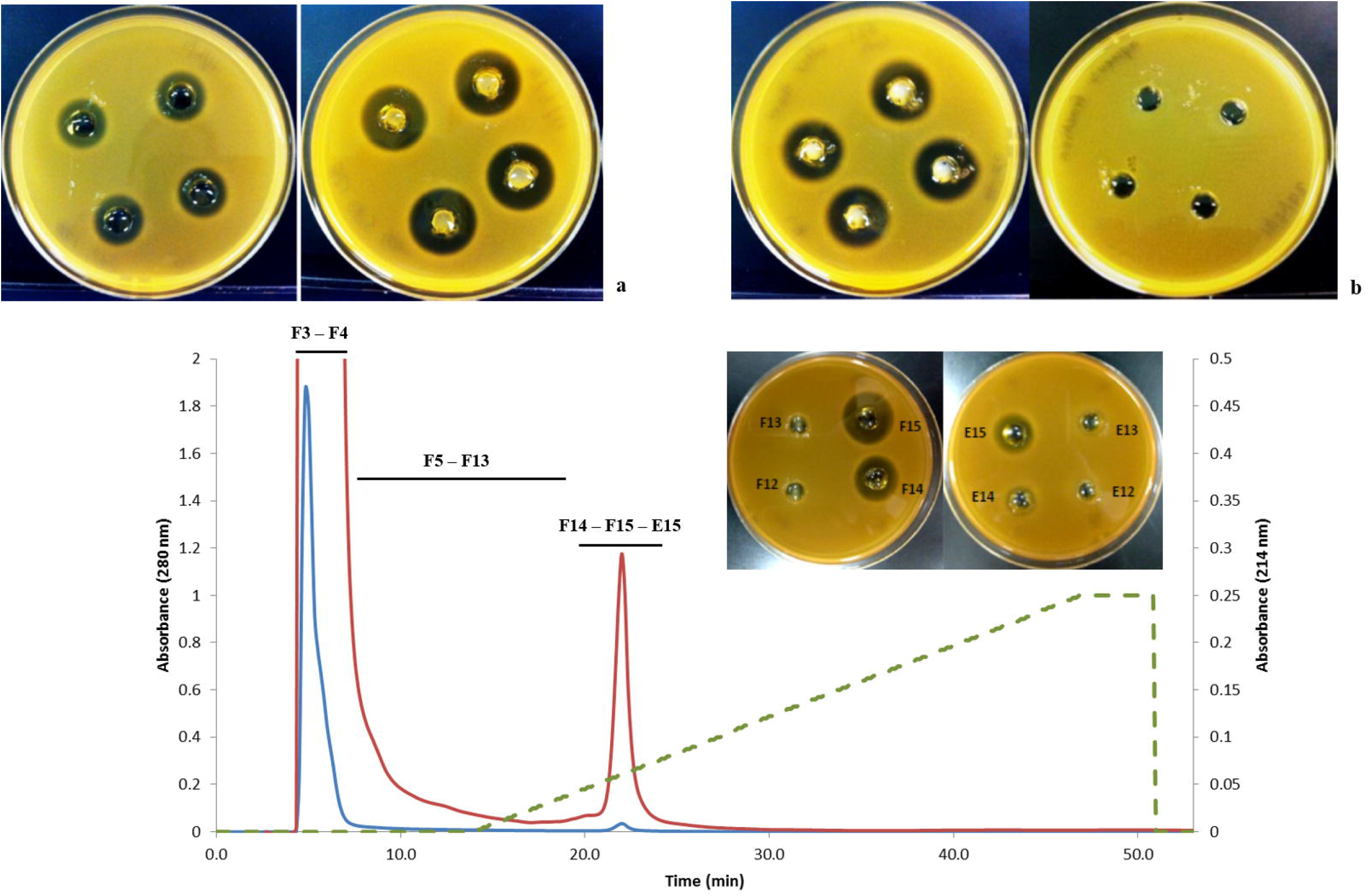
RP-HPLC elution profile of the *L. plantarum* B21 FPLC purified bacteriocin. Top figure A_220_; Bottom figure A_280_. An analytical ZORBAX Eclipse C_18_ column fitted to an Agilent 1100 RP-HPLC equipment was used with a flow rate of 1mL/min. The bacteriocin was eluted with approximately 80 % water-ACN.

### Bacteriocin peptide *de novo* sequencing

Initial attempts to determine the N-terminal amino acid sequence of the purified *L. plantarum* B21 bacteriocin by Edman-degradation failed (data not shown), suggesting that the peptide may be N-terminally blocked. A *de novo* sequencing approach was then taken to identify the peptide sequence of the purified bacteriocin peptide using an ESI-LC-MS/MS technique and identified a strong precursor ion at m/z 1417.0630 (monoisotopic peak) which corresponded to an intact monoisotopic mass of 5664.252 (Fig 5a). This was consistent with the previous mass measurements from MALDI-TOF MS Tris-Tricine-SDS-PAGE. To improve the MS/MS data, a targeted run was setup in which the m/z 1417 precursor (with an isolation width of 4 Da) was subject to both CID and HCD at three different energy levels and activation Q of 0.25 (CID) or activation time of 0.1ms (HCD). The spectrum produced from HCD failed to identify any candidate sequence in the existing MASCOT databases, so the obtained sequence tags were extracted manually (PGWAVAAAGALG and AAVILGV, Fig 5b and 5c) and BLAST was used to search a six-frame translation of the *L. plantarum* B21 genome. These amino acid peptide sequences corresponded to a part of an open reading frame on a 20 kb native plasmid pB21AG01 (GenBank Accession No. CP025732). A putative bacteriocin mature protein sequence was obtained from the predicted ORF (Fig 5b). From the mass spectrum data it appeared that the N-terminal sequence of the peptide started with the sequence PGWAVAAAGALG (Fig 5b) and the AAVILGV (Fig 5c) sequence was close to the C-Terminus; however comparison to the predicted amino acid from the genome data, indicated that AAVILGV sequence occurs just before the PGWAVAAAGALG sequence in the predicted ORF and was not separated by more than ~4000 Da of mass (Fig 6). The precursor mass of the peptide was found to be 18Da less than the apparent mass of the predicted mature amino acid sequence. This result could be explained if the bacteriocin is a cyclic peptide. The formation of a peptide bond between the N and C-Terminus and resulting loss of water resulted in mass discrepancy of 18 Da between the gene sequence (5682 Da) and the experimental mass of 5664 Da. After the full putative peptide sequence was deduced from the genome data, the remainder of the peptide sequence could be traced and confirmed in the MS/MS data (Fig 5b and 5c) and the *L. plantarum* B21 circular bacteriocin peptide was named plantacyclin B21AG.

**Fig 5.**
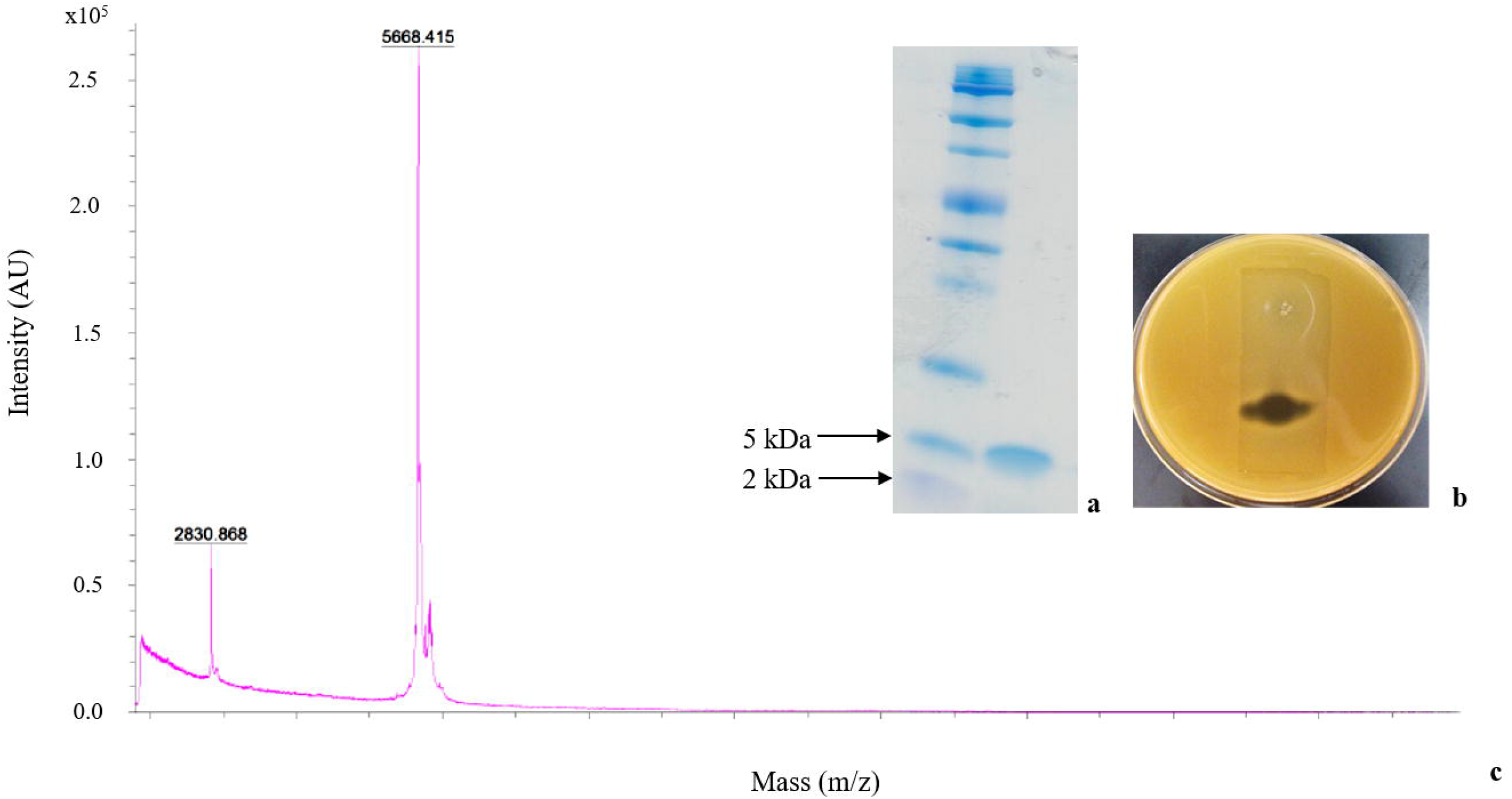
*De novo* peptide sequencing of the *L. plantarum* B21 bacteriocin using LC-MS/MS techniques. (a) The +4 species centred at 1417.81 was manually selected for fragmentation analysis. (b) Identification of the b ions. Peptide sequence manually extracted from the MS/MS data was PGWAVAAAGALG. (c) Identification of the late b ions. Peptide sequence obtained was AAVILGV.

**Fig 6.**
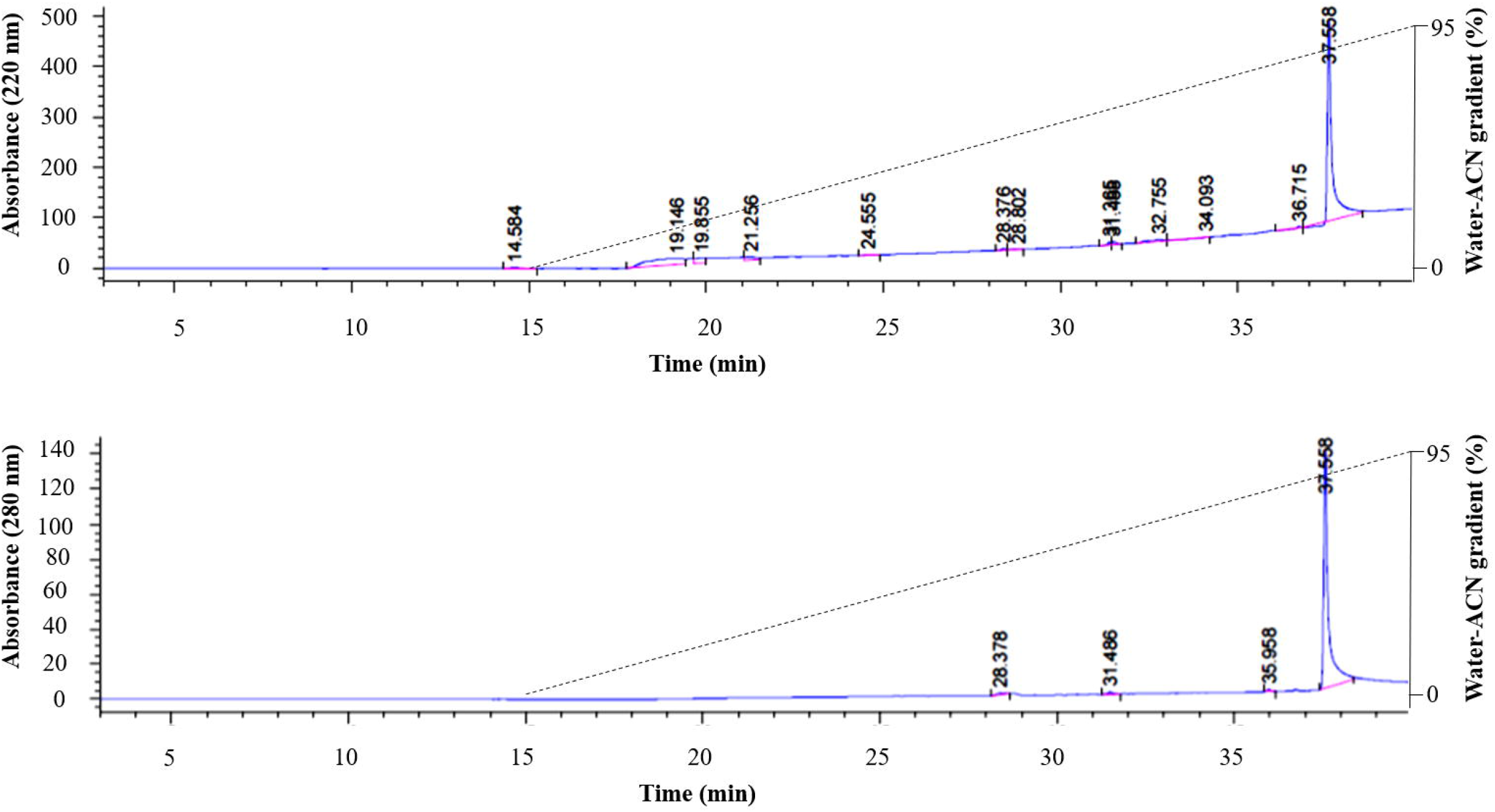
Determination of the amino acid and nucleotide sequence of plantacyclin B21AG. (a) Amino acid sequences obtained from *de novo* peptide sequencing using LC-MS/MS. (b) Plantacyclin B21AG mature peptide corresponding to the linear isotopic mass of 5682 Da, confirming the data obtained by MS/MS analysis. (c) Plantacyclin B21AG full putative sequence deduced from translation of the predicted ORF in pB21AG01. The hypothetical cleavage site of the leader peptide is indicated by a blue arrow. (d) The nucleotide sequence obtained from the *L. plantarum* B21 genome data corresponding to the plantacyclin B21AG amino acid sequence. The symbol (-) demonstrates the translation stop codon. (e) Predicted circular structure of plantacyclin B21AG. The black arrows show the main ion fragmentation site identified by the proteomic analysis.

The observed behaviour of plantacyclin B21AG in this study was consistent with the properties of a circular bacteriocin. Firstly, plantacyclin B21AG could not be sequenced by N-terminal sequencing, similar to other circular bacteriocins, including enterocin 62-6 [59], lactocyclicin Q [60] and butyrivibriocin AR10 [61]. Secondly, plantacyclin B21AG showed some level of resistance to proteolytic enzymes. It has been reported that circular bacteriocins are generally more stable than classical bacteriocins [59, 62, 63]. They are less susceptible to digestion by endoproteinases and this probably increases their spectrum of activity [60, 64]. Maqueda et al. (2004) [63] suggested that substantial part of the increased stability of enterocin AS-48 was due to the entropic constraints induced by the cyclic nature of the polypeptide chain. Plantacyclin B21AG was also shown to behave more hydrophobically on RP-HPLC column, requiring a high concentration (~80%) of ACN for elution. This behaviour has been observed for other circular bacteriocins like enterocin AS-48 [63] and enterocin 62-6 [59]. Finally, the protein sequence of the ORF corresponding to plantacyclin B21AG was shown to have similarity to other known circular bacteriocins.

### DNA and protein sequence analysis of plantacyclin B21AG

The putative bacteriocin peptide has 91 amino acids (pre-peptide) consisting of a 33-amino-acid leader peptide and a 58-amino-acid pro-peptide (Fig 6d). The putative cleavage site for removing the leader peptide is located between asparagine and isoleucine, and has homology to the leader peptide cleavage site in gassericin A and acidocin B [65, 66]. The 58-amino-acid peptide is predicted to undergo a post-translationally modification that results in the linking of the N-terminal asparagine to the C-terminal isoleucine, with the elimination of a water molecule, resulting in the active and mature B21AG molecule that is actually secreted. The leader peptides in circular bacteriocins vary in length, from 3 to more than 30 amino acids [21, 64, 67]. BLASTp analysis of the mature bacteriocin peptide (58 amino acids) showed 86% and 67% identity to the circular bacteriocin plantaricyclin A [32] produced by *L. plantarum* NI326 (accession number WP_053266997.1) and predicted circular bacteriocin pentocin KCA1 (accession number EIW14922), respectively. It also showed 65% identity to known circular bacteriocins produced by *Lactobacillus* species gassericin A [31] and acidocin B [68] (accession numbers WP_012621083.1 and CAA84399.1). These results demonstrate that plantacyclin B21AG is a new member of circular bacteriocin class produced by *Lactobacillus* spp. and it resembles a high similarity to the circular bacteriocin produced by the same species *L. plantarum*.

The amino acid composition of the mature circular bacteriocin consists of a very high proportion (59%) of hydrophobic amino acid residues (Ala, Val, Leu, Ile, Phe, Trp and Pro) and also uncharged hydrophilic amino acid residues (32%) (Gly, Ser, Thr and Gln). There is also high ratio of basic (Lys, Arg and His) relative to acidic amino acids (Asp) suggesting a strong basic protein character. It has been shown that the basic residues such as Lys present a highly localised positive charge on the surface of the circular bacteriocins structure which is responsible for attracting the peptide to the surface of the negatively charged membrane [64, 69]. Plantacyclin B21AG does not contain any cysteine residues, a characteristic which is shared by all known circular bacteriocins. A lack of cysteine pairs means it is the C-N terminal ligation, not disulphide bridges which are responsible for the unique characteristics of circular bacteriocins. This is in contrast to cyclotides, circularised bacteriocins found in plants which [70] contain disulphide bridges. The bacteriocin peptide also contains three aromatic residues (two Trp and one Phe).

### Genetic analysis of plantacyclin B21AG gene cluster

The complete genome sequencing data was used to determine the genes responsible for plantacyclin B21AG production and immunity (GenBank Accession No. CP010528). The *b21ag* gene (ORF-A) (Fig 7) responsible for bacteriocin production is located on a 20 kb native plasmid pB21AG01 (GenBank Accession No. CP025732). Apart from the structural gene, we identified six additional ORFs (Fig 7, ORF-B to ORF-G) that appeared to be involved in bacteriocin production and immunity with the bacteriocin gene cluster (Fig 7). Six out of seven genes have the same orientation (ORF A to F), while one remaining gene (ORF-G) has an opposite orientation. Homologues of ORF-G are found in the same orientation as the rest of the genes in the acidocin B and gassericin A gene clusters, indicating a rearrangement event occurred sometime within the *b21ag* gene cluster.

Presence of a putative immunity gene indicates why antimicrobial activity against closely-related *Lactobacillus* species was so strong. It is likely that expression of plantacyclin B21AG will be active against the producing cell, so an immunity gene is also expressed to prevent self-killing. Any closely-related Lactobacillus will therefore also be susceptible. Given the high temporal stability of plantacyclin B21AG which is almost surely more stable than any immunity proteins, this helps ensure continued selection for the bacteriocin gene cluster on the pB21AG01 plasmid, functioning partly as a selfish genetic element [71].

**Fig 7.**
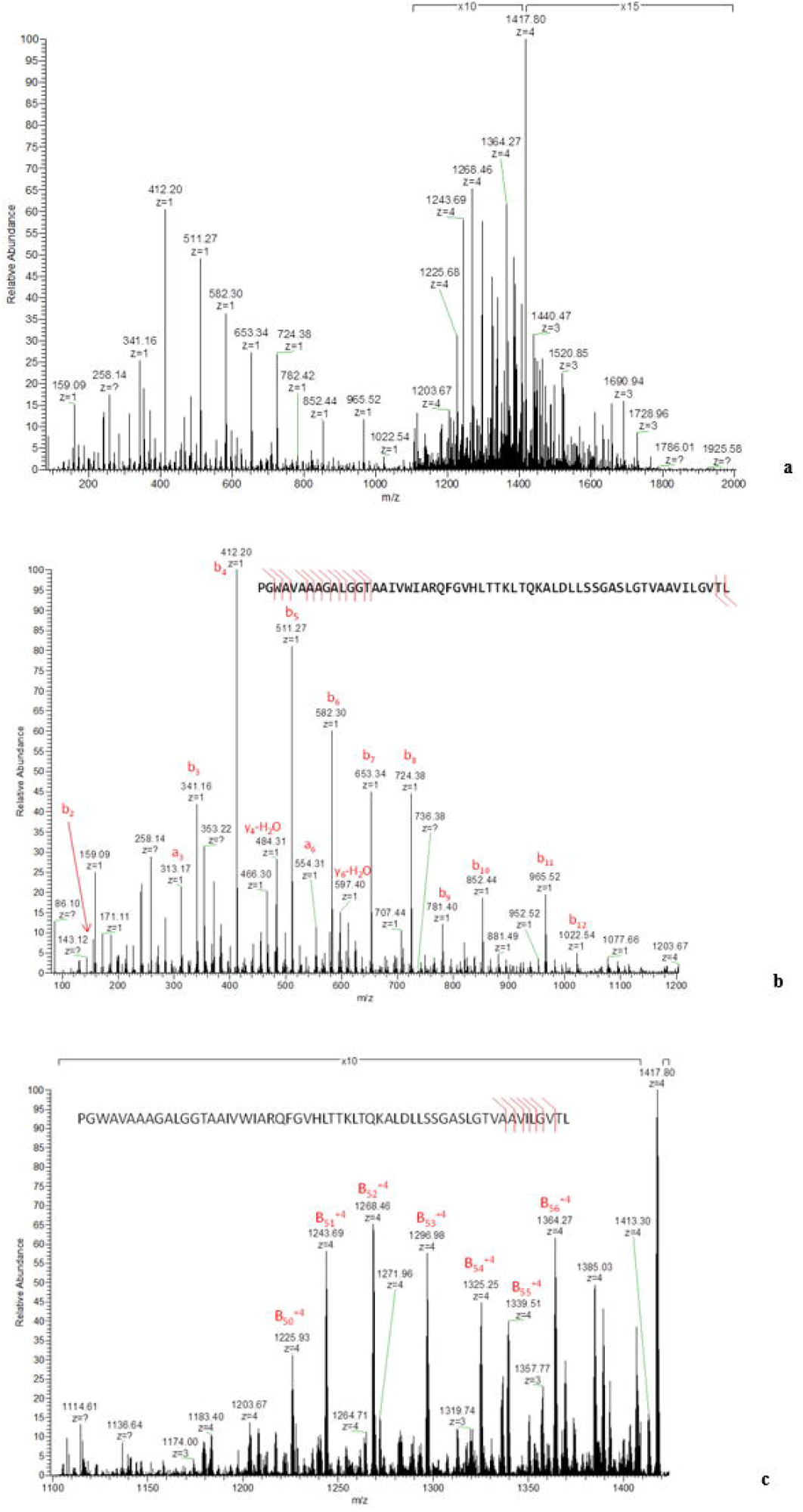
Schematic representation of the gene clusters involved in the production of circular bacteriocin gassericin A, acidocin B, pentocin KCA1, plantaricyclin A and plantacyclin B21AG. The genes are colour coded according to their known or putative functions. Green: Bacteriocin precursor; Blue: DUF95 family membrane protein; Yellow: Immunity protein; Orange: ABC transporter; Red: Membrane transporter; Grey: Unknown protein.

BLASTp analysis of the ORF-B putative protein sequence (157 amino acids) detected a putative conserved domain from the *DUF95* superfamily [72]. The ORF-B putative protein shows 94% sequence identity to *plcD*, which is a plantaricyclin A immunity protein [73]. It also shows some homology to other known circular bacteriocins, i.e 39%, 34%, 33% and 30% sequence identity to *PenD* from *L. pentosus* KCA1 [74], *AciD* from *L. acidophilus* M46 [53], *GaaD* from *L. gasseri* LA39 [75] and *BviE* from *Butyrivibrio fibrisolvens* AR10 [76], respectively. AciD belongs to the DUF95 family of membrane protein [53]. It has been demonstrated to be possibly involved in the biosynthesis of circular bacteriocin, as an immunity-associated transporter and as a secretion-aiding agent [77]. Kalmokoff, Cyr (76)] suggested that the *bviE* gene is involved in self-immunity to the circular bacteriocin, butyrivibriocin AR10. The dedicated immunity gene always appears to be located within the bacteriocin cluster [78]. The analysis of ORF-C putative protein sequence (54 amino acids) showed 89% identity to *plcI,* a plantaricyclin A putative immunity protein from *L. plantarum* NI326 [73]. It also shares 38% sequence identity with putative immunity proteins *GaaI* from *L. gasseri* LA39 [75] *and aciI* from *L. acidophilus* M46 [53] [79].

The ORF-D putative protein sequence (220 amino acids) shares 95% sequence identity to *plcT*, a plantaricyclin A ATP-binding ABC transporter protein [73]. It is also 49% identical to *penT* from *L. pentosus* KCA1 [74], and 46% identical to both *aciT* from *L. acidophilus* M46 [53] and *gaaT* from *L. gasseri* LA39 [75], which is involved in the transportation of gassericin A. It has been reported that dedicated transmembrane translocators belonging to the ATP-binding cassette (ABC) transporter superfamily are involved in the cleavage of the leader-peptide from Class II bacteriocins and in the transportation of the mature bacteriocin molecule across the cytoplasmic membrane [78]. These results suggested that ORF-D gene appeared be involved in the cleavage of the leader peptide and transport of plantacyclin B21AG bacteriocin protein. Similarly, ORF-E putative protein sequence (214 amino acids) showed 94% sequence identity to *plcE*, which is a plantaricyclin A membrane transporter [73]. It also showed 39%, 38% and 37% sequence identity to *penE* [74], *gaaE* [75] and *aciE* [53], respectively, encoding an ABC-2 transporter permease.

ORF-F putative protein sequence (173 amino acids) showed 90% sequence identity to plcB, which is a plantaricyclin A-related protein [73]. It is 31% and 30% identical to *penB* [74], and *aciB* [53], respectively. It also shares 30% sequence identity to *gaaB* [75], a membrane protein with 5 predicted transmembrane segments (TMS) from *L. gasseri* which is involved in production of the circular bacteriocin, gassericin A. Similarly, ORF-G showed 90% sequence identity to plcC, a plantaricyclin A related protein [73]. It also shares 30% sequence identity to *gaaC* [75] and *aciC* [53]. However, the locations of both ORF-F and ORF-G in the gene cluster of plantacyclin B21AG are different compared to gassericin A and acidocin B gene clusters as it is found downstream in the opposite orientation (Fig 7). Given the conserved nature of this gene in circular bacteriocin gene clusters and the fact that we have identified a putative promoter upstream of ORF-G, it is likely essential it is for bacteriocin production or immunity. If it were lost due to a recombination event, the gene cluster and corresponding immunity would cease to function and plantacyclin B21AG would likely kill the producer cell. However only rearrangements where ORF-G was retained with a new promoter has been found.

### Phylogenetic analysis of plantacyclin B21AG protein and sequence alignment

Phylogenetic analysis of the mature bacteriocin sequence of plantacyclin B21AG was performed against the mature sequences of experimentally-confirmed circular bacteriocins (Data S1), indicating two main groups of circular bacteriocins as previously described (Fig 8) [80]. Plantacyclin B21AG sits within the subgroup ii cluster, with other circular bacteriocin-producing *Lactobacillus* species and is closely related to plantaricyclin A. All of the mature sequences in subgroup ii contain 58 residues, whereas the length in subgroup i ranges from 60 – 70 residues.

**Fig 8.**
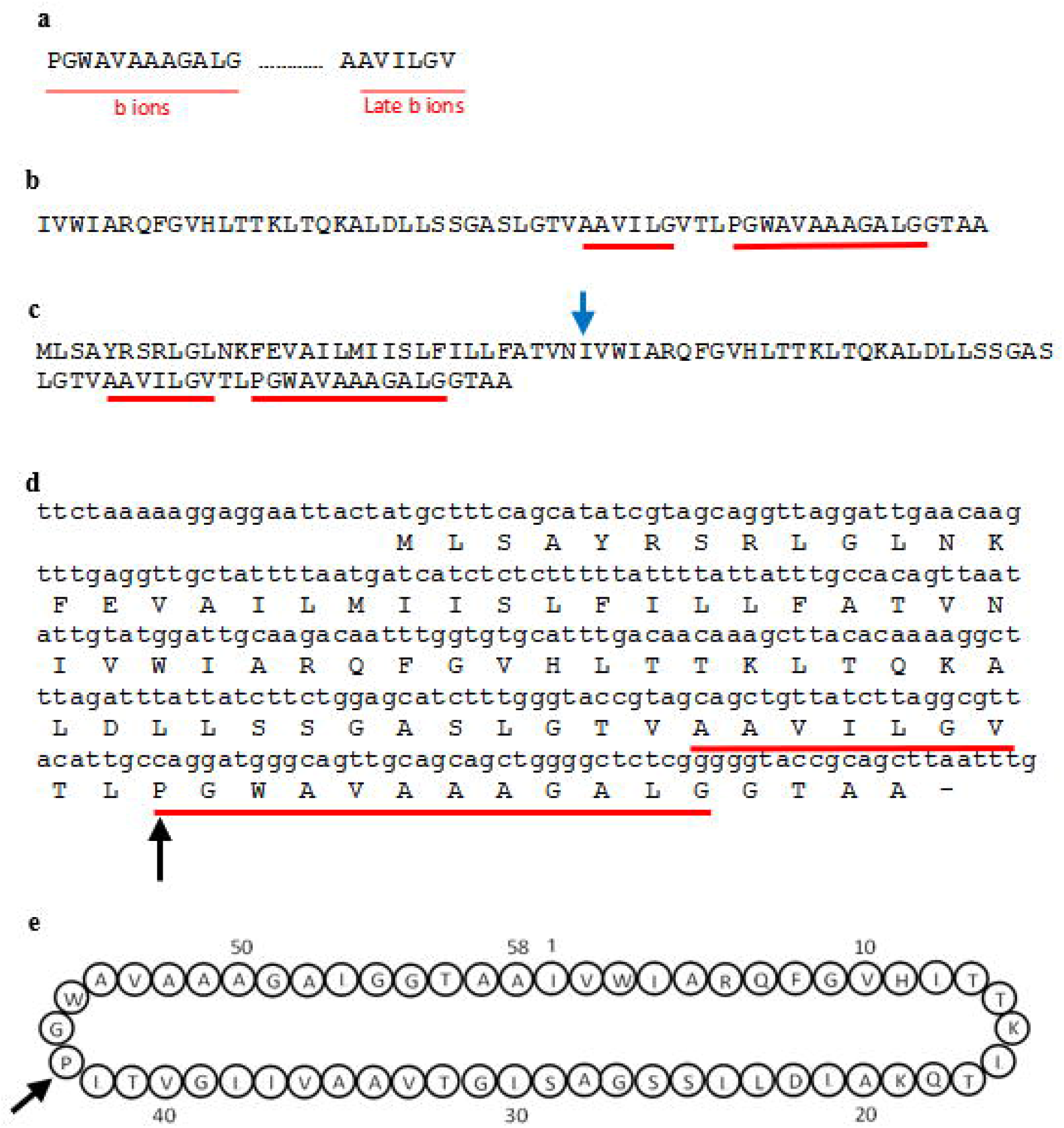
Phylogeny of the mature sequences of all known circular bacteriocins. The bacteriocins cluster into two main groups. Plantacyclin B21AG clusters within subgroup ii. Subgroup i contains the most variation of circular bacteriocins. Bootstrap values are shown. Asterisk shows plantacyclin B21AG.

## Conclusions

The antimicrobial agent produced by *L. plantarum* B21 was identified as a circular bacteriocin plantacyclin B21AG, corresponding to a molecular weight of 5668 Da, confirmed with MALDI-TOF MS and *de novo* sequencing using ESI-LC-MS/MS. The purification process demonstrated a yield of approximately 19% and the overall purification factor was more than 8,000-fold. A simple back calculation suggested *L. plantarum* B21 produces approximately 1 μg/mL bacteriocin in its CFS. The purified plantacyclin B21AG was shown to be temporally stable, thermostable, active across a wide pH range and partially resistant to proteolysis. Its spectrum of antimicrobial activity included killing activity against closely-related *Lactobacillus* species, as well as foodborne pathogens *Clostridium perfringens* and *Listeria monocytogenes*. This newly identified circular bacteriocin has many characteristics which would be desirable for food preservation

## Materials and methods

### Bacterial strains and culture conditions

The bacteriocin-producing organism, *L. plantarum* B21 [36] and the indicator strain, *L. plantarum* A6 [37], were taken from the RMIT University Culture Collection [38]. The remaining indicator strains, *L. plantarum* ATCC 8014, *L. arabinosus* 17-5, *L. brevis* 19012, *Lactococcus lactis* 345-18, *Listeria monocytogenes* 192/1-2 ACM 3173 and *Clostridium perfringens* 52/6-1 were purchased from the American Type Culture Collection (ATCC) or obtained from RMIT University Culture Collection. All LAB strains were grown in MRS broth (Oxoid, England) at 30°C for 24 hours without shaking. For growth of the LAB strains on solid media, MRS agar (Oxoid, England) was used and the plates were incubated at 30°C for 48 hours. *C. perfringens* and *L. monocytogenes* were grown anaerobically in Brain-heart infusion (BHI) (Oxoid, England) broth at 37°C and 30°C, respectively.

### Antimicrobial activity assay

The antimicrobial activity of *L. plantarum* B21 was examined using a well diffusion assay (WDA) [39]. Briefly, the cell free supernatant (CFS) of *L. plantarum* B21 was prepared from a 2% (v/v) inoculum of an overnight culture. Semi-solid MRS medium supplemented with 0.8% agar was mixed with approximately 10^6^ CFU/mL of the indicator strain and allowed to solidify. Wells of 8-mm in diameter were perforated and 100 μL of the CFS was placed into each well. The agar plates were held at 4°C for 2 h, followed by incubation at 30°C for 18-24 hours. The antimicrobial activities of the CFS were determined by measuring the diameters of the inhibition zones. Inhibition was recorded as negative if no zone was observed around the agar well. For quantitative measurement, two-fold serial dilutions of the CFS were prepared using PBS and the antimicrobial activities were expressed as arbitrary units (AU/mL) as described previously [40].

### Sensitivity to proteolytic enzymes

The CFS of *L. plantarum* B21 was harvested by centrifugation at 5,000 × g for 20 min at 4°C. The CFS was then neutralised to pH6.5 and treated separately with proteinase K, trypsin and pepsin (Sigma-Aldrich, USA) at a final concentration of 1mg/mL. Untreated CFS was included as a control whereas CFS treated with catalase (Sigma-Aldrich, USA) was used as a negative control. This was followed by WDA assay to examine the loss of antimicrobial activity. The diameters of the inhibition zones observed from the treated CFS were compared against those produced by untreated CFS or catalase-treated, used as positive and negative controls, respectively. The loss or reduction in the size of the inhibition zone was taken as an indication of the sensitivity of the antimicrobial activity to proteolytic digestion.

### Purification of bacteriocin

The *L. plantarum* B21 bacteriocin was purified from the CFS by a five-step protocol. CFS was harvested and, concentrated, followed by extraction, then parsed through size exclusion/desalting/buffer-exchange and finally processed through cation exchange chromatography by Fast Protein Liquid Chromatography (FPLC). All purification steps were carried out at room temperature. *L. plantarum* B21 was used as an inoculum 2% (v/v) into 165 mL of MRS broth and incubated at 30°C for 24 h without shaking. The CFS was prepared as previously described. The CFS was concentrated to 70 mL through a 10 kDa polyethersulfone membrane discs (Generon, UK) using an Amicon UF cell (Model 8200, Millipore, USA). The concentrated CFS was subjected to n-butanol extraction according to Abo-Amer [41] with a few modifications. The concentrated *L. plantarum* B21 CFS was extracted twice in 1:1 volume of water saturated n-butanol and centrifuged at 10,000 × g for 10 min. The n-butanol fraction containing the bacteriocin was freeze dried (FDU-8612, Operon Co. Ltd, Korea) to remove the solvent. The dried n-butanol fraction was redissolved in 20 mM sodium phosphate buffer, pH 6.0. Both the n-butanol and aqueous fractions were examined for antimicrobial activity. The redissolved protein sample was desalted using a NAP10 desalting column pre-packed with Sephadex G-25 resin (NAP™-10 Columns, Sephadex™ G25, GE Healthcare Life Sciences, UK) and eluted in 20 mM sodium phosphate buffer (pH 6.0). Purification of the desalted bacteriocin protein was performed using an Uno S-6 prepacked monolith cation exchange column (12×55 mm, Bio-Rad, USA) on a FPLC system (BioLogic DuoFlow System, Bio-Rad, USA). The Uno S-6 column was equilibrated with 20 mM sodium phosphate buffer, pH 6.0 (buffer A) and the desalted bacteriocin protein was eluted from the column with a linear NaCl gradient (0-1 M) in buffer A. A total of 33 fractions of 3 mL were collected and assayed for the antimicrobial activity. The two FPLC-fractions showing the strongest bacteriocin activity (F14 and F15) were pooled together (6 mL) and concentrated/buffer exchanged using an Amicon^®^ Ultra-4 Centrifugal Filter Units (3 kDa, Millipore, Ireland). The final volume of the purified bacteriocin protein fraction was reduced to 200 μL. The concentration of the bacteriocin was measured after each step of protein purification using Pierce™ BCA Protein Assay Kit (Thermo Fisher Scientific, USA).

### Determination of molecular weight and purity

The molecular weight and purity of the FPLC-purified bacteriocin was determined using tris-tricine-sodium dodecyl sulfate-polyacrylamide gel electrophoresis (Tris-Tricine-SDS-PAGE, 16.5% resolving gel) as previously described [42]. After electrophoresis at 60 V for 30 min followed by 100 V for 1.5 h, half of the gel was stained using a Coomassie-based staining solution (Expedeon, UK), while the other half was fixed in a fixing solution containing 20% isopropanol and 10% acetic acid for the examination of antimicrobial activity as described previously [43]. *L. plantarum* A6 was used an indicator strain. Precision Plus Protein™ Dual Xtra Standards (Bio-Rad, USA) was used to estimate the size of the bacteriocin. The same sample was also investigated by matrix-assisted laser desorption/ionization time-of-flight mass spectrometry (MALDI-TOF MS) [44] using an Autoflex Speed MALDI-TOF instrument (Bruker, Germany) containing a 355-nm Smartbeam II laser for desorption and ionization. The acceleration and reflector voltages were 20 and 23.4 kV in pulsed ion extraction mode. A molecular mass gate of 450 Da improved the measurement by filtering out most matrix ions. To determine the purity of the sample, the bacteriocin was diluted 10-fold in 0.1% trifluoroacetic acid (TFA) and subjected to analytical reverse-phase high-performance liquid chromatography (RP-HPLC) [45] using an Agilent 1100 HPLC system fitted with a ZORBAX Eclipse XDB analytical C_18_ column (4.6×150 mm, 5 μm particle size, Agilent Technologies, USA). The bacteriocin protein was eluted using a 40-min linear water-ACN gradient (increasing the gradient to 95%).

### *De novo* LC-MS/MS peptide sequencing

A *de novo* sequencing approach was used to identify the peptide sequence of the purified bacteriocin peptide using an ESI-LC-MS/MS technique at the Bio21 institute proteomics facility (University of Melbourne, Australia). *L. plantarum* B21 purified bacteriocin was analysed using an LTQ™ Orbitrap Elite ETD (Thermo Scientific, USA) coupled to an UltiMate 3000 RSLCnano System (Dionex, Thermo Scientific, USA). The nanoLC system was equipped with an Acclaim™ Pepmap™ 100 C_18_ nano-trap column (1×2 cm, 5 μm particle size, Thermo Scientific, USA) and an Acclaim™ Pepmap™ 100 C_18_ analytical column (length 15 cm, 5 μm particle size, Thermo Scientific, USA). 2 μL of the bacteriocin sample was loaded onto the trap column at 3% (v/v) ACN (Merck, USA) containing 0.1% (v/v) formic acid (Sigma-Aldrich, USA) for 5 min before the enrichment column was switched in-line with the analytical column. An initial run on the LTQ™ Orbitrap Elite ETD mass spectrometer (Thermo Scientific, USA) was set to operate in data-dependent mode, where subsequent MS/MS spectra was acquired for the top five peaks by collision induced dissociation (CID) activation, followed by higher-energy collision dissociation (HCD). To improve the MS/MS information a targeted run was setup where the precursor (with an isolation width of 4 Da) was subject to both CID and HCD at three different energy levels (CID: 26, 32, 36, and HCD: 24, 26, 30) and activation Q of 0.25 (CID) or activation time of 0.1 ms (HCD). The resulting fragment ion spectra were recorded in the Orbitrap at a resolution of 240,000. The spectrum produced from HCD at 26 eV was submitted to MASCOT.

### Genetic analysis of bacteriocin gene cluster

Sequence tags were extracted manually from the spectrum resulting from the de novo LC-MS/MS peptide sequencing and BLAST against the six-frame translation of the total *L. plantarum* B21 Illumina genome data (GenBank Accession No. CP010528). The DNA sequence of the bacteriocin was found to be located on a native plasmid designated as pB21AG01 (GenBank Accession No. CP025732). Plasmid annotation was performed using Rapid Annotation using Subsystem Technology (RAST) (http://rast.nmpdr.org/) [46] to identify the gene cluster involved in the production of the bacteriocin. All open reading frames (ORFs) identified were then searched against NCBI database using BLAST (http://www.ncbi.nlm.nih.gov) [47] to identify genes associated with bacteriocin production. Plantacyclin B21AG was aligned against other experimentally-confirmed circular bacteriocins using Clustal Omega [48] Clustal Omega (https://www.ebi.ac.uk/Tools/msa/clustalo/) and exported to fasta format, which was coverted to phylip format using Sequence Conversion (http://sequenceconversion.bugaco.com/converter/biology/sequences/fasta_to_phylip.php). This was used as input for RAxML (raxmlHPC-PTHREADS-SSE3 version 8.2.10) [49] using the following parameters for ML + rapid bootstrap analysis with 100 replicates: -T 2 -f a -x 285 -m PROTGAMMABLOSUM62 -p 639 -N 100

The bipartitions output file was used in FigTree version 1.4.4 (http://tree.bio.ed.ac.uk/software/figtree/) for viewing/manipulation.

## Acknowledgements

This work was supported by RMIT University. We are grateful to Mr. Frank Antolasic for his technical assistance in MALDI-TOF-MS analysis.

## Author Contributions

Conceived and designed the experiments: ATS. Provided bacteria strain: BKM. Performed the experiments: AG TTHV MCG. Analysed the data: AG TTHV MCG BV. Wrote manuscript: AG MCG BV. Editing: ATS MCG BV

## Notes

### Competing Interest Statement

The authors have declared no competing interest.

## References

1. Parada JL, Caron CR, Medeiros ABP, Soccol CR. Bacteriocins from lactic acid bacteria: purification, properties and use as biopreservatives. Brazilian Archives of Biology and Technology. 2007;50(3):512–42.

2. Kirk M FL, Glass K, Hall G. Foodborne Illness, Australia, Circa 2000 and Circa 2010.. Emerging Infectious Diseases. 2014;20(11):1857–64. doi: https://dx.doi.org/10.3201/eid2011.131315.

3. Hall G, Kirk MD, Becker N, Gregory JE, Unicomb L, Millard G, et al. Estimating foodborne gastroenteritis, Australia. Emerging infectious diseases. 2005;11(8):1257.

4. Molognoni L, Daguer H, Motta GE, Merlo TC, Lindner JDD. Interactions of preservatives in meat processing: Formation of carcinogenic compounds, analytical methods, and inhibitory agents. Food Research International. 2019;125:108608. doi: https://doi.org/10.1016/j.foodres.2019.108608.

5. Yang S-C, Lin C-H, Sung CT, Fang J-Y. Antibacterial activities of bacteriocins: application in foods and pharmaceuticals. Frontiers in microbiology. 2014;5:241.

6. Fabiszewska AU, Zielińska KJ, Wróbel B. Trends in designing microbial silage quality by biotechnological methods using lactic acid bacteria inoculants: a minireview. World Journal of Microbiology and Biotechnology. 2019;35(5):76. doi: 10.1007/s11274-019-2649-2.

7. Fardet A, Rock E. In vitro and in vivo antioxidant potential of milks, yoghurts, fermented milks and cheeses: a narrative review of evidence. Nutrition Research Reviews. 2018;31(1):52–70. Epub 2017/10/02. doi: 10.1017/S0954422417000191.

8. Carr FJ, Chill D, Maida N. The lactic acid bacteria: a literature survey. Critical reviews in microbiology. 2002;28(4):281–370.

9. Von Wright A, Axelsson L. Lactic acid bacteria: an introduction. Lactic acid bacteria: Microbiological and functional aspects. 2011:1–17.

10. Sanlibaba P, Gucer Y. ANTIMICROBIAL ACTIVITY OF LACTIC ACID BACTERIA. Agriculture and Food. 2015.

11. Favaro L, Penna ALB, Todorov SD. Bacteriocinogenic LAB from cheeses-application in biopreservation? Trends in food science & technology. 2015;41(1):37–48.

12. Perez RH, Zendo T, Sonomoto K. Novel bacteriocins from lactic acid bacteria (LAB): various structures and applications. Microbial cell factories. 2014;13(1):1.

13. Nes IF, Yoon S, Diep DB. Ribosomally synthesiszed antimicrobial peptides (bacteriocins) in lactic acid bacteria: a review. Food Science and Biotechnology. 2007;16(5):675.

14. Klaenhammer TR. Genetics of bacteriocins produced by lactic acid bacteria. FEMS microbiology reviews. 1993;12(1-3):39–85.

15. Potter A, Ceotto H, Coelho MLV, Guimarães AJ, Bastos MdCdF. The gene cluster of aureocyclicin 4185: the first cyclic bacteriocin of Staphylococcus aureus. Microbiology. 2014;160(5):917–28. doi: doi:10.1099/mic.0.075689-0.

16. Himeno K, Rosengren KJ, Inoue T, Perez RH, Colgrave ML, Lee HS, et al. Identification, Characterization, and Three-Dimensional Structure of the Novel Circular Bacteriocin, Enterocin NKR-5-3B, from Enterococcus faecium. Biochemistry. 2015;54(31):4863–76. doi: 10.1021/acs.biochem.5b00196.

17. Scholz R, Vater J, Budiharjo A, Wang Z, He Y, Dietel K, et al. Amylocyclicin, a Novel Circular Bacteriocin Produced by &lt;span class=&quot;named-content genus-species&quot; id=&quot;named-content-1&quot;&gt;Bacillus amyloliquefaciens&lt;/span&gt; FZB42. Journal of Bacteriology. 2014;196(10):1842. doi: 10.1128/JB.01474-14.

18. Kurata A, Yamaguchi T, Kira M, Kishimoto N. Characterization and heterologous expression of an antimicrobial peptide from Bacillus amyloliquefaciens CMW1. Biotechnology & Biotechnological Equipment. 2019;33(1):886–93. doi: 10.1080/13102818.2019.1627246.

19. Samyn B, Martinez-Bueno M, Devreese B, Maqueda M, Gálvez A, Valdivia E, et al. The cyclic structure of the enterococcal peptide antibiotic AS-48. FEBS Letters. 1994;352(1):87–90. doi: 10.1016/0014-5793(94)00925-2.

20. Tomita H, Fujimoto S, Tanimoto K, Ike Y. Cloning and genetic and sequence analyses of the bacteriocin 21 determinant encoded on the Enterococcus faecalis pheromone-responsive conjugative plasmid pPD1. Journal of Bacteriology. 1997;179(24):7843. doi: 10.1128/jb.179.24.7843-7855.1997.

21. Martin-Visscher LA, van Belkum MJ, Garneau-Tsodikova S, Whittal RM, Zheng J, McMullen LM, et al. Isolation and characterization of carnocyclin a, a novel circular bacteriocin produced by *Carnobacterium maltaromaticum* UAL307. Applied and environmental microbiology. 2008;74(15):4756–63. doi: 10.1128/AEM.00817-08. PubMed PMID: 18552180.

22. Kemperman R, Kuipers A, Karsens H, Nauta A, Kuipers O, Kok J. Identification and Characterization of Two Novel Clostridial Bacteriocins, Circularin A and Closticin 574. Applied and Environmental Microbiology. 2003;69(3):1589. doi: 10.1128/AEM.69.3.1589-1597.2003.

23. Egan K. Discovery and evaluation of novel and characterised bacteriocins for future applications. Cork, Ireland: University College Cork; 2018.

24. Borrero J, Brede DA, Skaugen M, Diep DB, Herranz C, Nes IF, et al. Characterization of Garvicin ML, a Novel Circular Bacteriocin Produced by &lt;em&gt;Lactococcus garvieae&lt;/em&gt; DCC43, Isolated from Mallard Ducks (&lt;em&gt;Anas platyrhynchos&lt;/em&gt;). Applied and Environmental Microbiology. 2011;77(1):369. doi: 10.1128/AEM.01173-10.

25. Sawa N, Zendo T, Kiyofuji J, Fujita K, Himeno K, Nakayama J, et al. Identification and Characterization of Lactocyclicin Q, a Novel Cyclic Bacteriocin Produced by *Lactococcus* sp. Strain QU 12 Applied and Environmental Microbiology. 2009;75:1552–8.

26. Masuda Y, Ono H, Kitagawa H, Ito H, Mu F, Sawa N, et al. Identification and Characterization of Leucocyclicin Q, a Novel Cyclic Bacteriocin Produced by &lt;span class=&quot;named-content genus-species&quot; id=&quot;named-content-1&quot;&gt;Leuconostoc mesenteroides&lt;/span&gt; TK41401. Applied and Environmental Microbiology. 2011;77(22):8164. doi: 10.1128/AEM.06348-11.

27. Wirawan RE, Swanson KM, Kleffmann T, Jack RW, Tagg JR. Uberolysin: a novel cyclic bacteriocin produced by Streptococcus uberis. Microbiology. 2007; 153(5): 1619–30. doi: doi:10.1099/mic.0.2006/005967-0.

28. Acedo JZ, van Belkum MJ, Lohans CT, McKay RT, Miskolzie M, Vederas JC. Solution Structure of Acidocin B, a Circular Bacteriocin Produced by &lt;span class=&quot;named-content genus-species&quot; id=&quot;named-content-1&quot;&gt;Lactobacillus acidophilus&lt;/span&gt; M46. Applied and Environmental Microbiology. 2015;81(8):2910. doi: 10.1128/AEM.04265-14.

29. Kalmokoff ML, Cyr TD, Hefford MA, Whitford MF, Teather RM. Butyrivibriocin AR10, a new cyclic bacteriocin produced by the ruminal anaerobe Butyrivibrio fibrisolvens AR10: characterization of the gene and peptide. Canadian Journal of Microbiology. 2003;49(12):763–73. doi: 10.1139/w03-101.

30. Collins FWJ, O’Connor PM, O’Sullivan O, Gómez-Sala B, Rea MC, Hill C, et al. Bacteriocin Gene-Trait matching across the complete Lactobacillus Pan-genome. Scientific Reports. 2017;7(1):3481. doi: 10.1038/s41598-017-03339-y.

31. Kawai Y, Saito T, Kitazawa H, Itoh T. Gassericin A; an Uncommon Cyclic Bacteriocin Produced by *Lactobacillus gasseri* LA39 Linked at *N-* and *C*-Terminal Ends. Bioscience, Biotechnology, and Biochemistry. 1998;62(12):2438–40. doi: 10.1271/bbb.62.2438.

32. Borrero J, Kelly E, Connor PM, Kelleher P, Scully C, Cotter PD, et al. Plantaricyclin A, a Novel Circular Bacteriocin Produced by &lt;span class=&quot;named-content genus-species&quot; id=&quot;named-content-1&quot;&gt;Lactobacillus plantarum&lt;/span&gt; NI326: Purification, Characterization, and Heterologous Production. Applied and Environmental Microbiology. 2018;84(1):e01801–17. doi: 10.1128/AEM.01801-17.

33. Gabrielsen C, Brede DA, Nes IF, Diep DB. Circular bacteriocins: biosynthesis and mode of action. Applied and environmental microbiology. 2014;80(22):6854–62.

34. Tran KTM, May BK, Smooker PM, Van TTH, Coloe PJ. Distribution and genetic diversity of lactic acid bacteria from traditional fermented sausage. Food Research International. 2011;44(1):338–44.

35. Tran KTM. Investigation of The native microflora and isolation of protective starter culture for a traditional Vietnamese fermented meat. Melbourne: RMIT University; 2010.

36. Golneshin A, Adetutu E, Ball AS, May BK, Van TTH, Smith AT. Complete genome sequence of Lactobacillus plantarum strain B21, a bacteriocin-producing strain isolated from Vietnamese fermented sausage nem chua. Genome announcements. 2015;3(2):e00055–15.

37. Golneshin A, Gor M-C, Van TTH, May B, Moore RJ, Smith AT. Draft Genome Sequence of Lactobacillus plantarum Strain A6, a Strong Acid Producer Isolated from a Vietnamese Fermented Sausage (Nem Chua). Genome announcements. 2017;5(41).

38. Tran KT, May BK, Smooker PM, Van TT, Coloe PJ. Distribution and genetic diversity of lactic acid bacteria from traditional fermented sausage. Food Research International. 2011;44(1):338–44.

39. Tagg J, McGiven A. Assay system for bacteriocins. Applied microbiology. 1971;21(5):943.

40. Todorov SD. Bacteriocin production by Lactobacillus plantarum AMA-K isolated from Amasi, a Zimbabwean fermented milk product and study of the adsorption of bacteriocin AMA-K to Listeria sp. Brazilian Journal of Microbiology. 2008;39:178–87.

41. Abo-Amer AE. Characterization of a bacteriocin-like inhibitory substance produced by Lactobacillus plantarum isolated from Egyptian home-made yogurt. ScienceAsia. 2007;33(3):313–9.

42. Schägger H, Von Jagow G. Tricine-sodium dodecyl sulfate-polyacrylamide gel electrophoresis for the separation of proteins in the range from 1 to 100 kDa. Analytical biochemistry. 1987;166(2):368–79.

43. Van R, Dicks, Chikindas. Isolation, purification and partial characterization of plantaricin 423, a bacteriocin produced by Lactobacillus plantarum. Journal of Applied Microbiology. 1998;84(6): 1131–7. doi: 10.1046/j.1365-2672.1998.00451.x.

44. Vater J, Kablitz B, Wilde C, Franke P, Mehta N, Cameotra SS. Matrix-assisted laser desorption ionization-time of flight mass spectrometry of lipopeptide biosurfactants in whole cells and culture filtrates of Bacillus subtilis C-1 isolated from petroleum sludge. Applied and Environmental Microbiology. 2002;68(12):6210–9.

45. Elegado FB, Kim WJ, Kwon DY. Rapid purification, partial characterization, and antimicrobial spectrum of the bacteriocin, Pediocin AcM, from< i> Pediococcus acidilactici</i> M. International journal of food microbiology. 1997;37(1):1–11.

46. Aziz RK, Bartels D, Best AA, DeJongh M, Disz T, Edwards RA, et al. The RAST Server: rapid annotations using subsystems technology. BMC genomics. 2008;9(1):75.

47. Altschul SF, Madden TL, Schäffer AA, Zhang J, Zhang Z, Miller W, et al. Gapped BLAST and PSI-BLAST: a new generation of protein database search programs. Nucleic acids research. 1997;25(17):3389–402.

48. Madeira F, Park YM, Lee J, Buso N, Gur T, Madhusoodanan N, et al. The EMBL-EBI search and sequence analysis tools APIs in 2019. Nucleic Acids Res. 2019;47(W1):W636–W41. doi: 10.1093/nar/gkz268. PubMed PMID: 30976793.

49. Stamatakis A. RAxML version 8: a tool for phylogenetic analysis and post-analysis of large phylogenies. Bioinformatics. 2014;30(9):1312–3. doi: 10.1093/bioinformatics/btu033.

50. Omar NB, Abriouel H, Lucas R, Martínez-Cañamero M, Guyot J-P, Gálvez A. Isolation of bacteriocinogenic Lactobacillus plantarum strains from ben saalga, a traditional fermented gruel from Burkina Faso. International Journal of Food Microbiology. 2006;112(1):44–50. doi: 10.1016/j.ijfoodmicro.2006.06.014.

51. Garneau S, Ference CA, van Belkum MJ, Stiles ME, Vederas JC. Purification and characterization of brochocin A and brochocin B (10-43), a functional fragment generated by heterologous expression in Carnobacterium piscicola. Applied and environmental microbiology. 2003;69(3):1352–8.

52. Babasaki K, Takao T, Shimonishi Y, Kurahashi K. Subtilosin A, a new antibiotic peptide produced by Bacillus subtilis 168: isolation, structural analysis, and biogenesis. Journal of biochemistry. 1985;98(3):585–603.

53. Acedo JZ, van Belkum MJ, Lohans CT, McKay RT, Miskolzie M, Vederas JC. Solution structure of acidocin B, a circular bacteriocin produced by Lactobacillus acidophilus M46. Applied and environmental microbiology. 2015;81(8):2910–8.

54. González B, Arca P, Mayo B, Suárez JE. Detection, purification, and partial characterization of plantaricin C, a bacteriocin produced by a Lactobacillus plantarum strain of dairy origin. Applied and Environmental Microbiology. 1994;60(6):2158–63.

55. Morgan S, Ross RP, Hill C. Bacteriolytic activity caused by the presence of a novel lactococcal plasmid encoding lactococcins A, B, and M. Applied and Environmental Microbiology. 1995;61(8):2995–3001.

56. Kabuki T, Saito T, Kawai Y, Uemura J, Itoh T. Production, purification and characterization of reutericin 6, a bacteriocin with lytic activity produced by Lactobacillus reuteri LA6. International Journal of Food Microbiology. 1997;34(2):145–56. doi: https://doi.org/10.1016/S0168-1605(96)01180-4.

57. Maldonado A, Ruiz-Barba JL, Jiménez-Díaz R. Purification and Genetic Characterization of Plantaricin NC8, a Novel Coculture-Inducible Two-Peptide Bacteriocin from Lactobacillus plantarum NC8. Applied and Environmental Microbiology. 2003;69(1):383–9. doi: 10.1128/aem.69.1.383-389.2003.

58. Smaoui S, Elleuch L, Bejar W, Karray-Rebai I, Ayadi I, Jaouadi B, et al. Inhibition of fungi and gram-negative bacteria by bacteriocin BacTN635 produced by Lactobacillus plantarum sp. TN635. Applied biochemistry and biotechnology. 2010;162(4):1132–46.

59. Dezwaan DC, Mequio MJ, Littell JS, Allen JP, Rossbach S, Pybus V. Purification and characterization of enterocin 62-6, a two-peptide bacteriocin produced by a vaginal strain of Enterococcus faecium: Potential significance in bacterial vaginosis. Microbial ecology in health and disease. 2007;19(4):241–50.

60. Sawa N, Zendo T, Kiyofuji J, Fujita K, Himeno K, Nakayama J, et al. Identification and characterization of lactocyclicin Q, a novel cyclic bacteriocin produced by Lactococcus sp. strain QU 12. Applied and environmental microbiology. 2009;75(6):1552–8.

61. Kalmokoff M, Teather R. Isolation and characterization of a bacteriocin (Butyrivibriocin AR10) from the ruminal anaerobe Butyrivibrio fibrisolvens AR10: evidence in support of the widespread occurrence of bacteriocin-like activity among ruminal isolates of B. fibrisolvens. Applied and environmental microbiology. 1997;63(2):394–402.

62. Maqueda M, Sánchez□Hidalgo M, Fernández M, Montalbán□López M, Valdivia E, Martinez□Bueno M. Genetic features of circular bacteriocins produced by Gram□positive bacteria. FEMS microbiology reviews. 2008;32(1):2–22.

63. Maqueda M, Gálvez A, Bueno MM, Sanchez-Barrena MJ, González C, Albert A, et al. Peptide AS-48: prototype of a new class of cyclic bacteriocins. Current Protein and Peptide Science. 2004;5(5):399–416.

64. Borrero J, Brede DA, Skaugen M, Diep DB, Herranz C, Nes IF, et al. Characterization of garvicin ML, a novel circular bacteriocin produced by Lactococcus garvieae DCC43, isolated from mallard ducks (Anas platyrhynchos). Applied and environmental microbiology. 2011;77(1):369–73.

65. Kawai Y, Saito T, Kitazawa H, Itoh T. Gassericin A; an uncommon cyclic bacteriocin produced by Lactobacillus gasseri LA39 linked at N-and C-terminal ends. Bioscience, biotechnology, and biochemistry. 1998;62(12):2438–40.

66. Leer RJ, van der Vossen JM, van Giezen M, Van Noort Johannes M, Pouwels PH. Genetic analysis of acidocin B, a novel bacteriocin produced by Lactobacillus acidophilus. Microbiology. 1995;141(7):1629–35.

67. Martínez-Bueno M, Maqueda M, Gálvez A, Samyn B, Van Beeumen J, Coyette J, et al. Determination of the gene sequence and the molecular structure of the enterococcal peptide antibiotic AS- 48. Journal of Bacteriology. 1994;176(20):6334–9.

68. Leer RJ, van der Vossen JMBM, van Giezen M, van Noort Johannes M, Pouwels PH. Genetic analysis of acidocin B, a novel bacteriocin produced by Lactobacillus acidophilus. Microbiology. 1995;141(7):1629–35. doi: doi:10.1099/13500872-141-7-1629.

69. Martin-Visscher LA, Gong X, Duszyk M, Vederas JC. The three-dimensional structure of carnocyclin A reveals that many circular bacteriocins share a common structural motif. Journal of Biological Chemistry. 2009;284(42):28674–81.

70. Craik DJ, Daly NL, Bond T, Waine C. Plant cyclotides: A unique family of cyclic and knotted proteins that defines the cyclic cystine knot structural motif11Edited by P. E. Wright. Journal of Molecular Biology. 1999;294(5):1327–36. doi: https://doi.org/10.1006/jmbi.1999.3383.

71. Ito Y, Kawai Y, Arakawa K, Honme Y, Sasaki T, Saito T. Conjugative Plasmid from &lt;em&gt;Lactobacillus gasseri&lt;/em&gt; LA39 That Carries Genes for Production of and Immunity to the Circular Bacteriocin Gassericin A. Applied and Environmental Microbiology. 2009;75(19):6340. doi: 10.1128/AEM.00195-09.

72. Marchler-Bauer A, Bo Y, Han L, He J, Lanczycki CJ, Lu S, et al. CDD/SPARCLE: functional classification of proteins via subfamily domain architectures. Nucleic acids research. 2016;45(D1):D200–D3.

73. Borrero J, Kelly E, O’Connor PM, Kelleher P, Scully C, Cotter PD, et al. Purification, characterization and heterologous production of plantaricyclin A, a novel circular bacteriocin produced by Lactobacillus plantarum NI326. Applied and Environmental Microbiology. 2017:AEM. 01801–17.

74. Anukam KC. Pentocin KCA1: a novel circular bacteriocin gene encoded in the genome of Lactobacillus pentosus KCA1 with putative basic property. Journal of African Association of Physiological Sciences. 2015;2(2):117–29.

75. Kawai Y, Kusnadi J, Kemperman R, Kok J, Ito Y, Endo M, et al. DNA sequencing and homologous expression of a small peptide conferring immunity to gassericin A, a circular bacteriocin produced by Lactobacillus gasseri LA39. Applied and environmental microbiology. 2009;75(5): 1324–30.

76. Kalmokoff M, Cyr T, Hefford M, Whitford M, Teather R. Butyrivibriocin AR10, a new cyclic bacteriocin produced by the ruminal anaerobe Butyrivibrio fibrisolvens AR10: characterization of the gene and peptide. Canadian journal of microbiology. 2003;49(12):763–73.

77. Mu F, Masuda Y, Zendo T, Ono H, Kitagawa H, Ito H, et al. Biological function of a DUF95 superfamily protein involved in the biosynthesis of a circular bacteriocin, leucocyclicin Q. Journal of bioscience and bioengineering. 2014;117(2):158–64.

78. Nes IF, Diep DB, Håvarstein LS, Brurberg MB, Eijsink V, Holo H. Biosynthesis of bacteriocins in lactic acid bacteria. Antonie van Leeuwenhoek. 1996;70(2-4): 113–28.

79. Ito Y, Kawai Y, Arakawa K, Honme Y, Sasaki T, Saito T. Conjugative plasmid from Lactobacillus gasseri LA39 that carries genes for production of and immunity to the circular bacteriocin gassericin A. Applied and environmental microbiology. 2009;75(19):6340–51.

80. Gabrielsen C, Brede DA, Nes IF, Diep DB. Circular Bacteriocins: Biosynthesis and Mode of Action. Applied and Environmental Microbiology. 2014;80(22):6854. doi: 10.1128/AEM.02284-14.

